# External error attribution dampens efferent-based predictions but not proprioceptive changes in hand localization

**DOI:** 10.1101/2020.02.05.936062

**Authors:** Raphael Q. Gastrock, Shanaathanan Modchalingam, Bernard Marius ’t Hart, Denise Y. P. Henriques

**Affiliations:** Centre for Vision Research, York University, Toronto, Ontario, Canada, M3J 1P3; Department of Psychology, York University, Toronto, Ontario, Canada, M3J 1P3; School of Kinesiology and Health Science, York University, Toronto, Ontario, Canada, M3J 1P3

## Abstract

In learning and adapting movements in changing conditions, people attribute the errors they experience to a combined weighting of internal or external sources. As such, error attribution that places more weight on external sources should lead to decreased updates in our internal models for movement of the limb or estimating the position of the effector, i.e. there should be reduced implicit learning. However, measures of implicit learning are the same whether or not we induce explicit adaptation with instructions about the nature of the perturbation. Here we evoke clearly external errors by either demonstrating the rotation on every trial, or showing the hand itself throughout training. Implicit reach aftereffects persist, but are reduced in both groups. Only for the group viewing the hand, changes in hand position estimates suggest that predicted sensory consequences are not updated, but only rely on recalibrated proprioception. Our results show that estimating the position of the hand incorporates source attribution during motor learning, but recalibrated proprioception is an implicit process unaffected by external error attribution.

## Introduction

Knowing our limbs’ positions is crucial for our ability to move competently. Moreover, changing circumstances may cause movement errors, which require us to adapt our motor control to restore performance ^[1-5]^. When errors are not caused by our own motor system, but are instead externally caused, the way in which movements are adapted to counter them should change ^[6-10]^. Externally caused errors should also affect our estimate of the position of our limb, but this has not been directly investigated yet. Here, we introduce two types of movement feedback to investigate how our limb position estimates may be affected when errors are clearly not caused by the individual.

In reaching movements, adaptive changes that result from small or gradually introduced visual or mechanical perturbations are traditionally considered as largely implicit^[2,11]^. Implicit adaptation is manifested by reach aftereffects, persistent deviations in hand movements after perturbation removal, suggesting an internal representational remapping has occurred in the brain ^[5,11-12]^. Reach aftereffects also occur with larger and abruptly introduced perturbations, as well as when participants are made aware of the nature of the perturbation. In these cases, explicit processes account for a part of the resulting adaptive change ^[13-18]^. Thus, both explicit and implicit processes contribute to adaptation ^[19-22]^. Here, we first quantify implicit and explicit contributions to learning with responses to different visual manipulations. These manipulations differentially demonstrate the nature and source of errors experienced, thereby varying the extent of external error attribution.

Motor adaptation leads not only to changes in motor performance, but previous research has also found that adapting reach movements to visual or mechanical perturbations leads to changes in proprioceptive estimates of hand location^[23-25]^, even if the two perturbations likely have different underlying mechanisms ^[26-27]^. This proprioceptive recalibration emerges quickly^[28-29]^ and reflects about 20% of the visual misalignment of the hand ^[23-24]^. Recalibrated proprioception is also preserved in aging ^[30]^and in different perturbations (rotations and translations ^[23]^, force fields ^[31]^, gains ^[32]^, split-belt walking ^[33-34]^). In visuomotor rotations, it seems that a visuo-proprioceptive discrepancy is sufficient to drive proprioceptive recalibration, and leads to reach aftereffects that mimic this proprioceptive shift ^[29,35-37]^. Thus, proprioceptive recalibration is ubiquitous, and seems to contribute to motor performance.

Apart from afferent proprioceptive information, hand localization is also based on predicted sensory consequences of the movement, calculated by internal forward models that use an efference copy of the outgoing motor command ^[38-39]^. These efferent-based updates are considered a pre-requisite for implicit adaptation ^[3,40]^, and seem to contribute to reach aftereffects separately from recalibrated proprioception ^[29,35,41]^. Efferents and non-visual afferents should both be present when estimating hand location after self-generated ‘active’ movements, while robot-generated ‘passive’ movements should only allow afferent-based proprioceptive signals. Thus, active and passive movements assess the relative contributions of afferent and efferent signals to hand position estimates ^[18,32,41]^, which should both be implicit.

Since both contributions to hand location estimates should be implicit, they should be reduced or not occur when errors are attributed externally, as implicit learning is engaged less or not at all. In other words, given that the cursor in a visuomotor rotation task is considered a representation of the hand ^[42]^, it would be intuitive for people to not update estimates of their hand location, when it is clear that the error is being caused by an external source. However, modulating explicit knowledge about the nature of the perturbation, by providing instructions or increasing the perturbation size, does not affect persistent shifts in both proprioceptive recalibration and updating of predicted sensory consequences ^[18]^. In the current study, we instead investigate the effect of the external attribution of errors on both afferent and efferent-based changes. To do this, we vary the extent that people attribute the error they experience to a cursor representing their hand position, while holding a robot manipulandum and training in a visuomotor rotation task (Fig. 1a-1d). The experiment consists of two sessions: a baseline, aligned session, where visual feedback of the cursor matched the actual hand position, and a rotated session where participants adapt to a 30° rotated hand-cursor (Fig. 2). In two groups that either receive instructions about the nature of the rotation and a strategy to counter for it, or not (Instructed and Control groups; Fig. 1a), we expect external error attribution to be minimal, as only explicit knowledge is modulated.

**Fig. 1:**
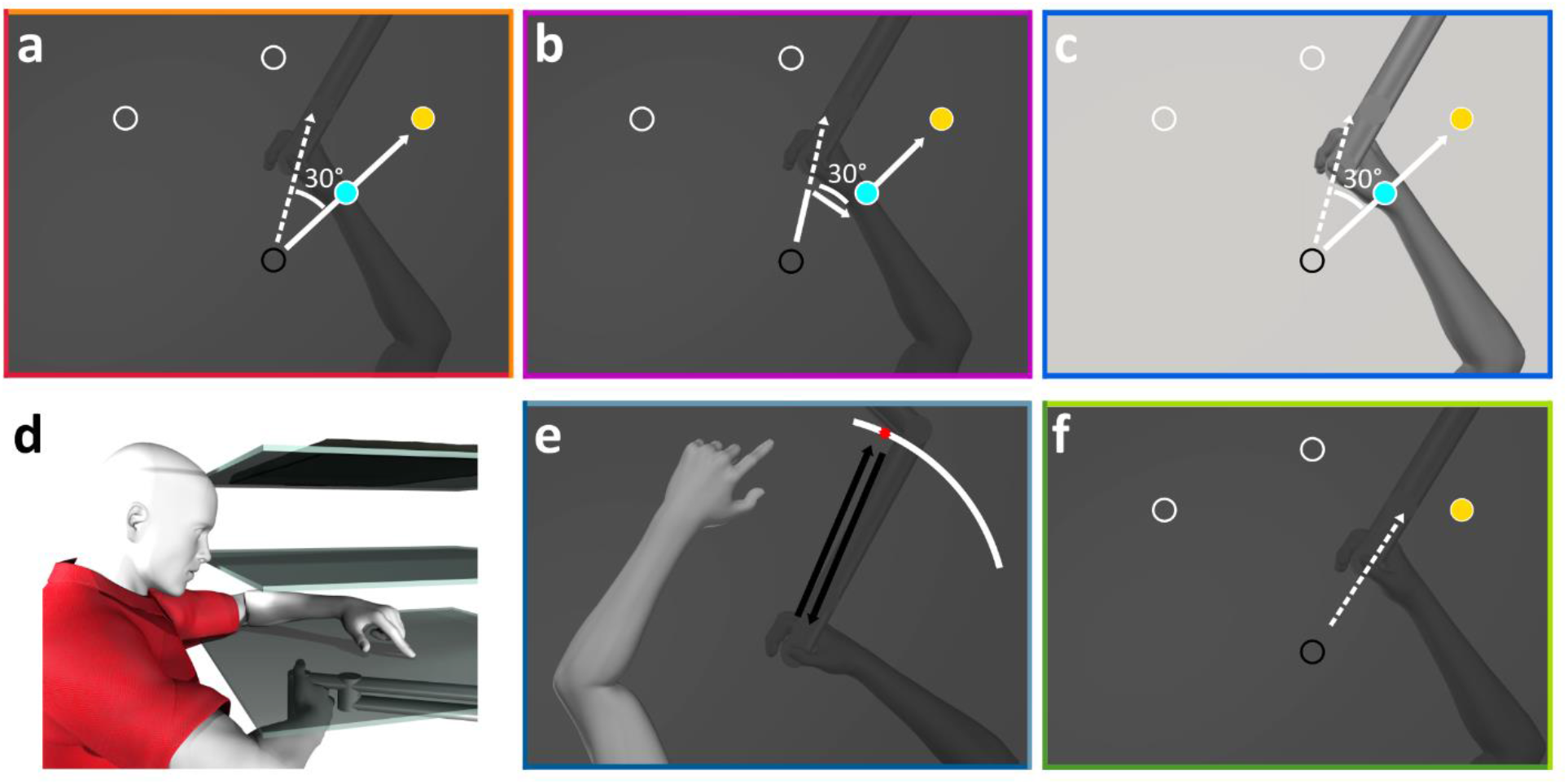
Experimental apparatus and stimuli. **a-c:** Top-down view displaying the different manipulations for the reach-training tasks, where the cursor (light blue) is rotated 30° CW. Reaches are made to one of three possible target locations (indicated as hollow white circles for reference), but only one target appeared on every trial (yellow disc). **a:** In both the Instructed and Control groups, participants do not see their hand, and the cursor has a constant rotation throughout each trial. **b:** Participants in the Cursor Jump group see the cursor “jump” to the 30° CW rotation mid-reach on every trial. **c:** In the Hand View group, participants see both their actual, illuminated hand and the cursor. **d:** Participants sit on an adjustable chair in a dark room and hold a robot manipulandum located below a touch screen (bottom surface), while viewing stimuli through a reflective tint (middle surface) which projects stimuli generated from a downward facing computer screen (top surface). **e:** Active and Passive Localization trials: Participants use their visible left hand to indicate on the touch screen where they have crossed the arc with their unseen right hand, after voluntarily generating a right-handed movement (active) or after a robot-generated movement (passive). **f:** No-cursor trials: Reaches are made to the same three targets in the absence of visual feedback of the cursor or hand.

**Fig. 2:**
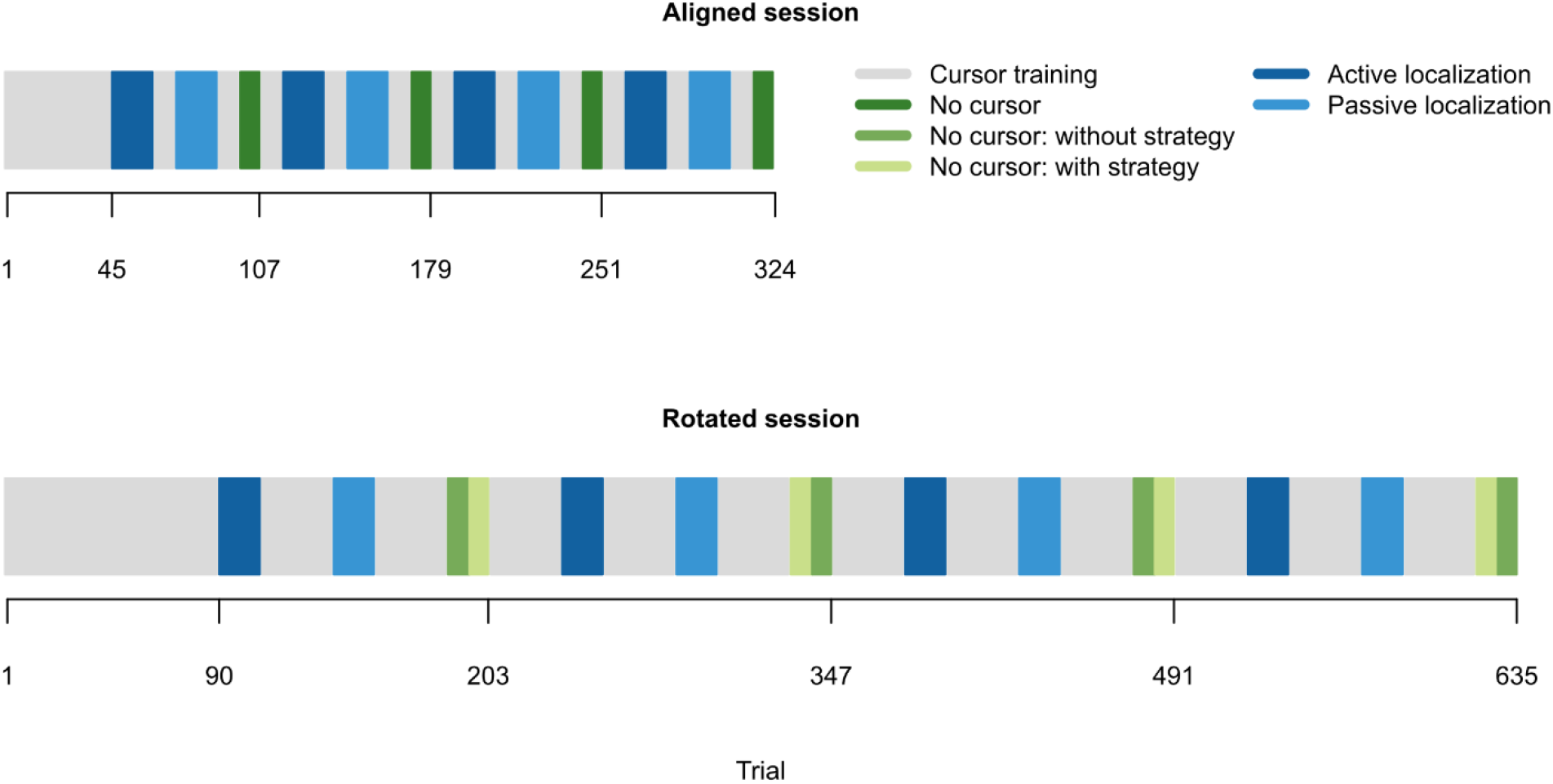
Experiment Schedule. **Top:** First session, and considered as baseline, where the cursor was aligned with the position of the right hand. Participants performed 45 cursor training trials followed by blocks of active localization (18 trials/block), passive localization (18 trials/block), and no-cursor trials (9 trials/block). Top-up cursor training trials (9 trials/block) were interleaved in between localization and no-cursor blocks. **Bottom:** Second session where the cursor was rotated 30° CW, relative to the position of the right hand. Participants performed 90 cursor training trials followed by blocks of active localization (18 trials/block), passive localization (18 trials/block), and two variations of no-cursor trials (with- or without-strategy; 9 trials/block). Top-up cursor training trials (30 trials/block) were interleaved in between localization and no cursor blocks. For both aligned and rotated sessions, passive localization always proceeded after active localization, as endpoint locations of the robot-generated movements in passive localization are based on locations that participants voluntarily moved towards during active localization. For no-cursor trials in the rotated session, the two variations are counterbalanced both within and between participants. That is, with- and without-strategy trials alternate within one participant, and the variation that an individual starts with is also alternated between participants.

In addition, we test two other groups that also do not receive instructions but either have visual feedback of the hand-cursor jump to the imposed rotation mid-reach on every training trial (Cursor Jump group; Fig. 1b) or a view of the actual hand of the participant is present along with the rotated cursor (Hand View group; Fig. 1c). We expect that these manipulations should make clear to participants that the cursor errors are caused externally. We interleave a localization task (Fig. 1e) and no-cursor reaches (Fig. 1f) across blocks of cursor training in both aligned and rotated sessions, to investigate how our manipulations affect changes in hand location estimates and motor behaviour respectively, following adaptation (Fig. 2). We hypothesize that with increased external error attribution, both changes in motor behaviour and shifts in afferent and efferent-based estimates of hand localization will decrease.

## Results

Before investigating how external error attribution affects changes in motor behaviour and hand localization, we first confirm that all groups appropriately counter the perturbation by the end of 90 training trials (Fig. 3a) and observe that reach trajectories are not qualitatively different (Fig. 4). We test for group differences at different time points during adaptation training (three blocks: trials 1-3, 4-6, 76-90) using a 3X4 mixed design ANOVA, with block (blocks 1, 2, and 3) as a within-subject factor and group (Control, Instructed, Cursor Jump, Hand View) as a between-subject factor. We find main effects of group (*F*(_3,86)_ = 5.678, *p* =.001, generalized eta squared (η^2^_G_) =.092, BF_incl_ > 1 · 10^6^) and block (*F*_(2,172)_ = 78.411, *p* <.001, η^2^_G_ =.307, BFincl > 3 · 10^14^), and a group X block interaction (*F*^(6,172)^ = 7.856, *p* <.001, η^2^_G_ =.118, BFincl > 4 · 10^5^). This suggests that, as expected, group differences in learning rates are modulated by the block of trials. Follow-up tests comparing each group to the Control group, show the expected initial advantage of instructions in reducing reach direction error within block one (Fig. 3a-3b), as only the Instructed group differs from the Control group (*t*_(148)_ = 4.632, *p* <.001, eta squared (η^2^) =.127, BF_10_ > 1 · 10^5^). In the second block (Fig. 3c), no groups differ from the Control group (Instructed: *t*_(148)_ = 1.922, *p* =.295, η^2^ =.024, BF_10_ = 6.506; Cursor Jump: *t*_(148)_ = 2.538, *p* =.071, η^2^ =.042, BF_10_ = 3.386; Hand View: *t*_(148)_ = 0.910, *p* =.934, η^2^ =.006, BF_10_ = 0.381). Bayesian analysis show moderate evidence for a difference between the Control group and the Instructed or Cursor Jump groups, but we note that these are calculated without correcting for multiplicity. For the last block (Fig. 3a,3d), an ANOVA on the effect of group on angular reach deviations shows that the groups do not differ from each other (*F*_(3,86)_ = 0.561, *p* =.642, η^2^_G_ =.019, BF_10_ = 0.115), suggesting that our manipulations do not affect the asymptotic level of adaptation. Thus, any effects of training on changes in motor behaviour and hand localization can’t be explained by levels of adaptation in the different groups.

**Fig. 3:**
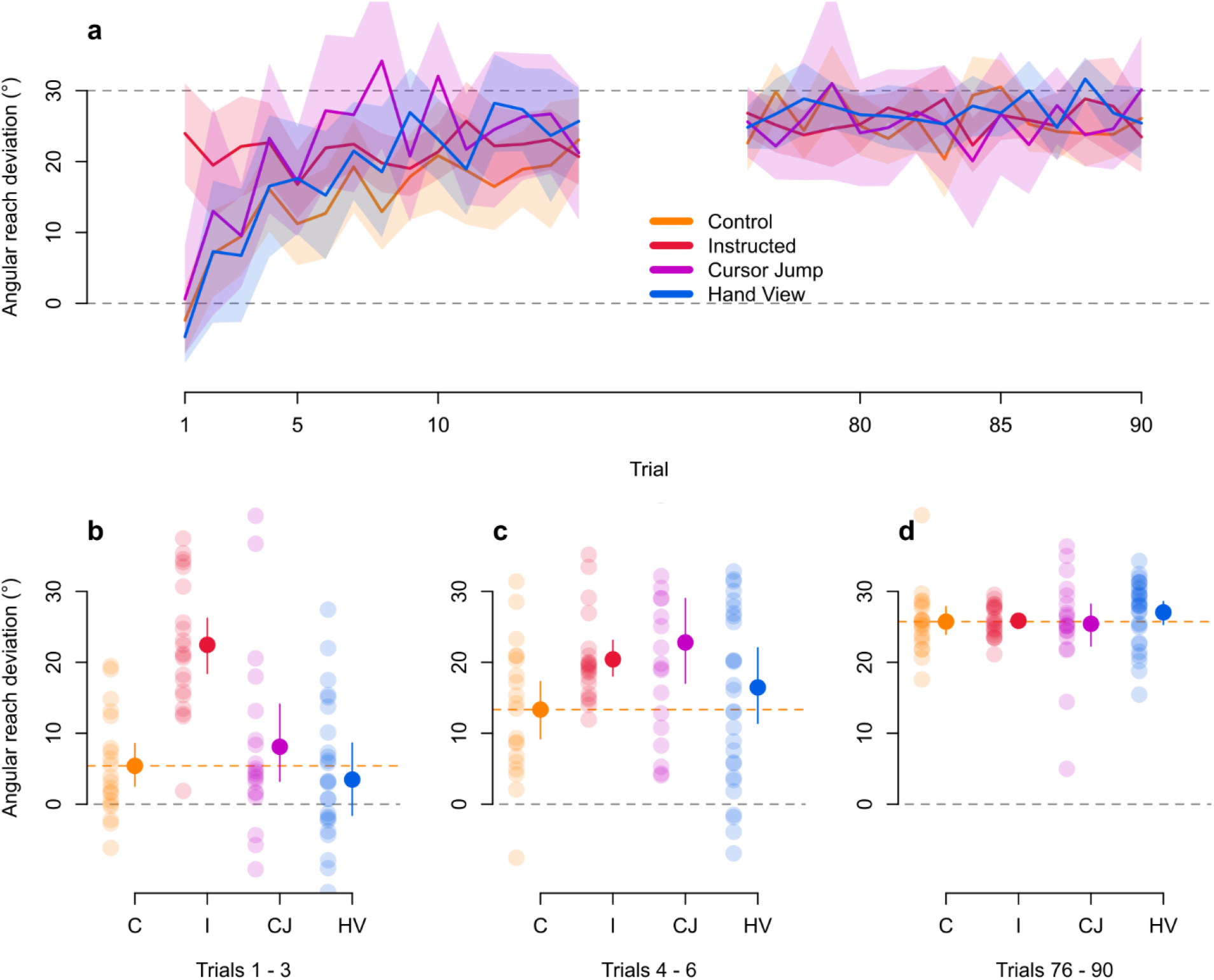
Rate of learning during adaptation training. **a:** Only the first and last 15 trials of adaptation training are shown. Grey dashed line at the 30° mark indicates the direction that the hand must deviate in order to fully and successfully counter for the perturbation. The grey dashed line at the 0° mark indicates reach directions similar to those in the baseline, aligned session (i.e., no compensation). The Instructed group shows an initial advantage in successfully countering for the perturbation as early as the first trial. There are no differences in reaches performed by participants from all groups for the last 15 trials. Solid lines are group means and shaded regions are corresponding 95% confidence intervals. **b-d:** Individual participant data from each group are shown, separated in three blocks of trial sets during adaptation training. Orange dashed line indicates mean for the Control group. Solid dots and error bars correspond to the group mean and bootstrapped 95% confidence intervals.

**Fig. 4:**
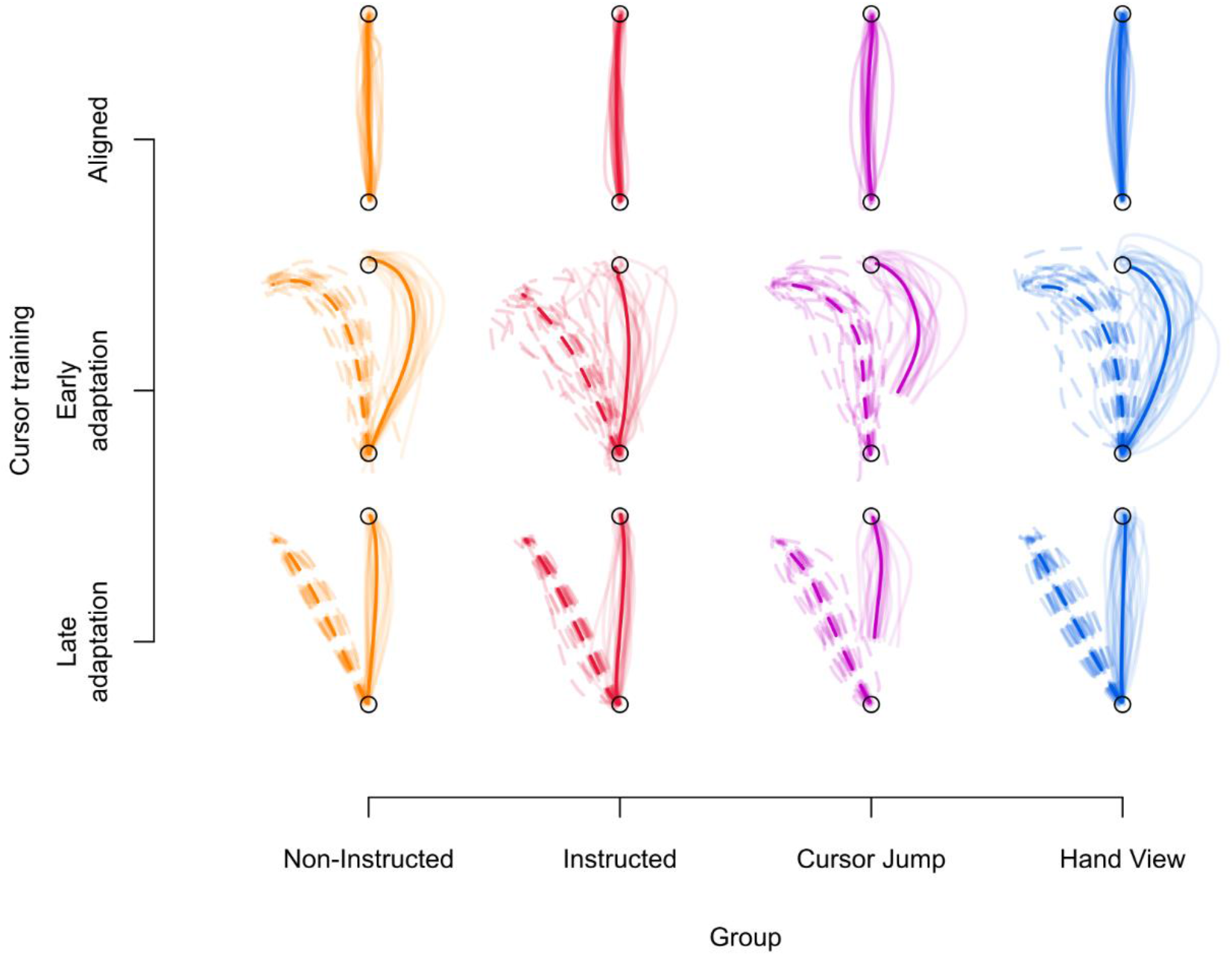
Individual and average reach trajectories. The trajectory of reaches across all participants within their respective groups are shown with light coloured lines. Each participant’s trajectory combines reaches during the last three trials of the first block of cursor training in the aligned session (**top**), as well as the first (**middle**) and last three (**bottom**) trials of the first block of cursor training in the rotated session. Light solid lines indicate the trajectory of the hand-cursor and light dashed lines indicate the trajectory of the hand. Only solid lines are shown in the aligned session, as both hand and hand-cursor trajectories are similar. Group means for the hand trajectories are indicated with the dark dashed line, and dark solid lines indicate the mean hand-cursor trajectory. All groups seem to produce similar reach trajectories, across the different time points in the experiment, regardless of condition. Moreover, despite curved reaches during early adaptation training, reach trajectories are straight towards the end of adaptation training.

### Implicit aftereffects persist despite external error attribution

To investigate the effects of external error attribution on changes in motor behaviour, we use no-cursor trials both before and after adaptation (Fig. 1f). After adaptation, however, we use a process dissociation procedure (PDP), a cognitive research methodology adapted by Werner et al. ^[16]^ for motor learning, which measures awareness by having participants either express or repress a learned movement (see also^[17-18,43]^). Here, we ask people to make open-loop reaches, and move their unseen right hand to targets, while either including any strategy they learned to counter for the perturbation (with-strategy reaches) or excluding it (without-strategy reaches). With explicit awareness about the nature of the perturbation, we expect a difference between these reaches, as the ability to consciously produce a strategy adds explicit contributions on top of implicit contributions to learning. Meanwhile, excluding a strategy reflects only implicit contributions, which are not consciously accessible. Thus, the PDP allows us to measure both implicit and explicit adaptation.

We first compare aligned no-cursor trials and without-strategy no-cursor reaches in the rotated session, to test for implicit reach aftereffects (Fig. 2, 5). We conduct a 2X4 mixed design ANOVA with session (aligned or rotated) as a within-subject factor and group as a between-subject factor. We confirm the presence of reach aftereffects with a main effect of session (*F*_(1,86)_ = 373.023, *p* <.001, η^2^_G_=.530, BF_incl_ = inf.). Moreover, we find a main effect of group (*F*_(3,86)_ = 16.576, *p* <.001, η^2^_G_=.230, BF_incl_ > 9 · 10^13^) and an interaction between session and group (*F*_(3,86)_ = 22.605, *p* <.001, η^2^_G_ =.170, BF_incl_ > 4 · 10^8^), suggesting that the effect of session is modulated by group. Follow-up tests show that aligned and without-strategy reach deviations differ within each group (Instructed: *t*_(86)_ = −11.830, *p* <.001, η^2^ =.619, BF_10_ > 1 · 10^6^; Control: *t*_(86)_ = −12.912, *p* <.001, η^2^ =.660, BF_10_ > 1 · 10^8^; Cursor Jump: *t*_(86)_ = −9.050, *p* <.001, η^2^ =.488, BF_10_ > 4 · 10^5^; Hand View: *t*_(86)_= −4.037, *p* <.001, η^2^ =.159, BF_10_ = 1 · 10^3^). This means that implicit reach aftereffects are present in each group. To address how the effect of session is modulated by group, follow-up tests compare implicit reach aftereffects for each group to those in the Control group. We find that the Instructed group doesn’t differ from the Control group (*t*_(86)_ = −0.722, *p* =.922, η^2^ =.006, odds = 0.099), but the Hand View (*t*_(86)_ = −7.538, *p* <.001, η_2_ =.398, odds > 7 · 10^3^) group does, suggesting that external error attribution in the Hand View group leads to reduced implicit reach aftereffects, compared to the Instructed and Control groups. Frequentist analysis shows that the Cursor Jump group differs from the Control group (*t*_(86)_ = −3.419, *p* =.004, η^2^ =.120), but this is not supported by Bayesian analysis (odds = 0.875). Furthermore, the reduction in aftereffects is more pronounced for the Hand View group compared to the Cursor Jump group (*t*_(86)_ = 3.818, *p* =.001, η^2^ =.145, odds = 2.220). In short, reach aftereffects persist across groups, but are greatly reduced for the Hand View group.

**Fig. 5:**
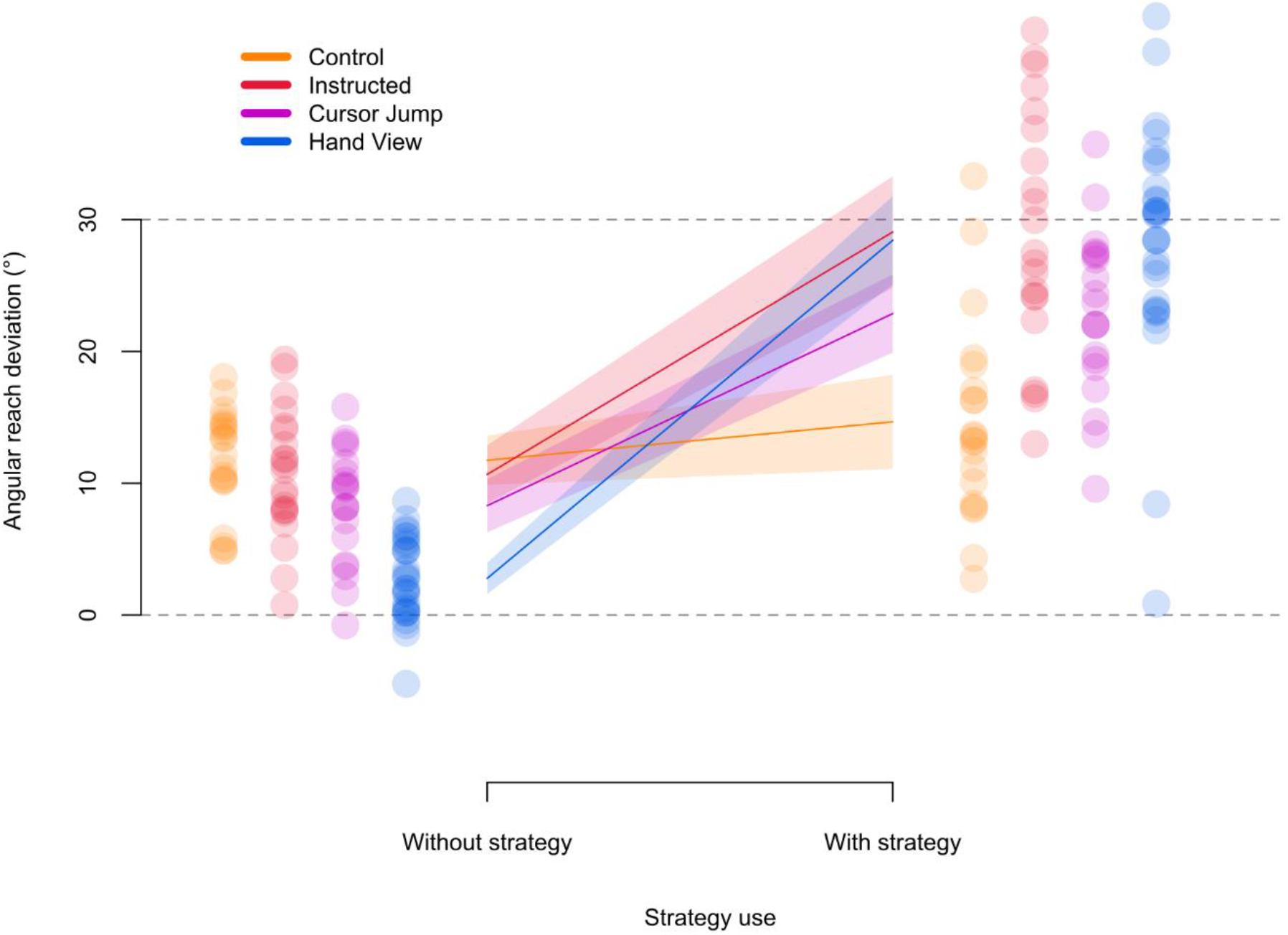
No cursor reaches and strategy use. Angular reach deviations of the hand per group, while either excluding (without-strategy) or including (with-strategy) any strategies developed during adaptation training. Grey dashed line at the 30° mark indicates angular reach deviations equivalent to full compensation for the perturbation, and grey dashed line at the 0° mark indicates reaches that did not correct for the perturbation. Only the Control group was unable to switch between excluding and including a strategy to counter for the perturbation. Implicit reach aftereffects, indicated by without-strategy angular reach deviations, are reduced for the Cursor Jump group and are further reduced in the Hand View group. Solid lines are group means and shaded regions are corresponding 95% confidence intervals. Individual participant data from each group are shown for both types of strategy use.

After confirming the presence of reach aftereffects, we use the PDP to assess explicit contributions to learning, by comparing with- and without-strategy no-cursor reaches (Fig. 5). We conduct a 2X4 mixed design ANOVA with strategy use (without-strategy or with-strategy) as a within-subject factor and group as a between-subject factor. We find main effects of strategy use (*F*_(1,86)_ = 285.493, *p* <.001, η^2^_G_ =.592, BF_incl_ = inf.) and group (*F*_(3,86)_ = 6.779, *p* <.001, η^2^_G_ =.118, BF_incl_ > 1 · 10^13^), and a strategy use and group interaction (*F*_(3,86)_ = 28.678, *p* <.001, η^2^_G_ =.304, BF_incl_ > 1 · 10^13^). This suggests that the effect of strategy use in at least one group is different from the other groups. Follow-up tests compare with- and without-strategy angular reach deviations for each group separately. We find no evidence for or against an effect of strategy use in the Control group (*t*_(86)_ = −1.529, *p* =.427, η^2^ =.026, BF_10_ = 0.940), but do see a difference in strategy use in the other groups (Instructed: *t*_(86)_ = −9.877, *p* <.001, η^2^ =.531, BF_10_ > 3 · 10^6^; Cursor Jump: *t*_(86)_ = −7.637, *p* <.001, η^2^ =.404, BF_10_> 5 · 10^4^; Hand View: *t*_(86)_ = −16.185, *p* <.001, η^2^ =.753, BF_10_ > 5 · 10^11^). Thus, despite receiving no instructions, both Cursor Jump and Hand View groups can evoke an explicit strategy like the Instructed group.

### Changes in afferent-based estimates of hand localization persist

We then investigate the effects of external error attribution on afferent and efferent-based shifts in hand location estimates. We use localization trials (Fig. 1e, 2), where participants indicate with their visible left hand, the position of their unseen right hand. Hand localization is either based on both afferent and efferent contributions (active localization) or based mainly on afferent contributions (passive localization). All groups appear to show shifts in hand localization, despite external error attribution (Fig. 6). Moreover, these shifts seem larger in active than passive localization for each group, except for the Hand View group (Fig. 6a-6b, 6d-6e). To test if training affected hand location estimates, we conduct a 2X2X4 mixed design ANOVA on localization error with session (aligned or rotated) and movement type (active or passive) as within-subject factors and group as a between-subject factor. We find a main effect of session (*F*_(1,86)_ = 82.972, *p* <.001, η^2^_G_ =.199, BF_incl_ = inf.) and group (*F*_(3,86)_ = 10.214, *p* <.001, η^2^_G_ =.195, BF_incl_ > 1 · 10^5^), an interaction between session and group (*F*_(3,86)_ = 2.895, *p* =.040, η^2^_G_ =.025, BF_incl_ = 354.651) and between session and movement type (*F*_(1,86)_ = 16.802, *p* <.001, η^2^_G_ =.004, BF_incl_ = 0.169). This suggests that estimates of hand position do shift despite external error attribution, but these shifts are modulated by group and movement type. Bayesian analysis suggests that including the session and movement type interaction does not lead to the best model (BF_10_best model > 1 · 10^34^).

**Fig. 6:**
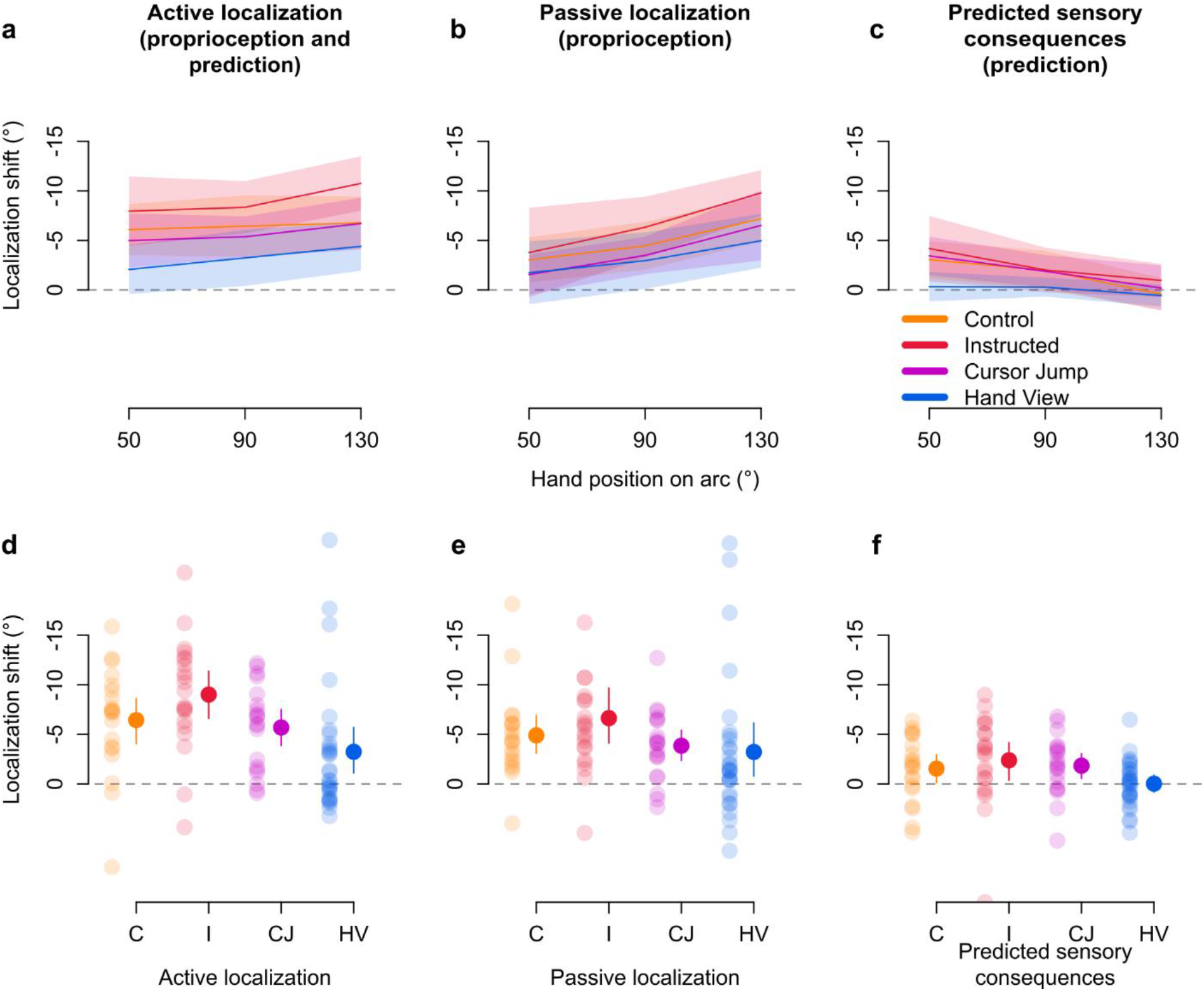
Afferent and efferent-based changes in hand location estimates. During localization trials, the arc stimulus is presented and participants either move, or are moved, towards different points on the arc. Shifts in localizing the unseen right hand following adaptation training after **a:** self-generated movements (active localization), **b:** robot-generated movements (passive localization), and **c:** the difference between active and passive localization as a measure of updates in efferent-based estimates (predicted sensory consequences). Grey dashed line at the 0° mark indicates the absence of shifts, while positive and negative values indicate the direction of shifts. Solid lines correspond to group means at each of three hand positions on the arc, which mark the position in polar coordinates of where the arc stimuli are centred on during these trials. These positions closely match the target locations during adaptation training and no-cursor reaches. Shaded regions are corresponding 95% confidence intervals. **d-f:** Individual participant data for shifts in hand localization are shown in transparent dots, separated according to group and movement type. Solid dots and error bars to the side of individual data correspond to group means and bootstrapped 95% confidence intervals.

Nonetheless, as planned, we consider movement type in the following frequentist test. We analyze the effects of group and movement type using a 2X4 mixed design ANOVA on localization shifts (i.e. difference in localization error between rotated and aligned sessions), with movement type as a within-subject factor and group as a between-subject factor. We find a main effect of movement type (*F*_(1,86)_ = 16.802, *p* <.001, η^2^_G_ =.016, BF_incl_ = 62.496) and group (*F*_(3,86)_ = 2.895, *p* =.040, η^2^_G_ =.085, BF_incl_= 2.540), but no interaction (*F*_(3,86)_ = 2.425, *p* =.071, η^2^_G_ =.007, BF_incl_ = 1.849), which is supported by

Bayesian analysis showing that the best model does not include this interaction (BF_10_ best model = 131.040). The main effect of movement type is expected because active movements contain afferent and efferent contributions to hand localization, while passive movements only have afferent contributions. For follow-up tests on the group effect, we compare the localization shifts of each group to the other groups regardless of movement type, and find that the Hand View group differs from the Instructed group (*t*_(86)_ = 2.901, *p* =.028, η^2^ =.089, odds = 14.120). Regardless, given the persistent shifts in hand position estimates, we investigate the afferent and efferent contributions for each group separately.

Passive localization should rely mainly on updated afferents, or recalibrated proprioception. We confirm the persistence of passive localization shifts across all groups with one-tailed t-tests that compare the mean passive localization shift of each group to zero (Instructed: *t*_(20)_ = −4.614, *p* <.001, *d* = 1.007, BF_10_= 348.746; Control: *t*_(19)_= −4.869, *p* <.001, *d* = 1.089, BF_10_ = 525.747; Cursor Jump: *t*_(19)_ = −4.832, *p* <.001, *d* = 1.080, BF_10_ = 488.283; Hand View: *t*_(28)_ = −2.372, *p* =.012, *d* = 0.440, BF_10_ = 4.201). These tests show that the attribution of error to external sources surprisingly does not reduce proprioceptive recalibration. Given that passive localization shifts reflect proprioceptive recalibration, a difference between active and passive localization shifts is likely due to efferent-based contributions. Thus, we measure efferent-based contributions or updates in predicted sensory consequences by removing afferent-based contributions (active minus passive; Fig. 6c,6f). We confirm the presence of updates in predictions for all groups with one-tailed t-tests comparing the mean shifts for each group to zero. We find that updates in predictions differ from zero for three groups (Instructed: *t*_(20)_ = −2.411, *p* =.013, *d* = 0.526, BF_10_= 4.570; Control: *t*_(19)_ = −2.101, *p* =.025, *d* = 0.470, BF_10_ = 2.729; Cursor Jump: *t*_(19)_ = −2.751, *p* =.006, *d* = 0.615, BF_10_ = 8.327), but not for the Hand View group (*t*_(28)_ = −0.037, *p* =.485, *d* = 0.007, BF_10_= 0.203). However, a Bayesian t-test comparing updates in predictions between the Control and Hand View groups provides little evidence for a difference between the two (BF_10_ = 1.225). On the other hand, reduced or absent updates in prediction could explain that active and passive localization shifts are not much different in the Hand View group. These results show that external error attribution might decrease or even eliminate efferent-based contributions to hand localization, but clearly does not affect afferent contributions to hand localization.

We then investigate whether the processes underlying afferent and efferent-based estimates of hand localization may independently be contributing to motor behaviour. Sensory prediction-error based learning should lead to updated predictions of hand location and contribute to reach aftereffects _[3,4,13,44-46]_, and aftereffects have been shown to emerge in the absence of updates to efferent-based predictions _[29,35-37]_, showing that recalibrated proprioception may be associated with both changes in hand location estimates and changes in behaviour _^[47]^_. When considering either passive localization shifts or updates in predictions and their respective relationships with angular reach deviations in without-strategy no-cursor trials (Fig. 7a-7b), we find that both share a small relationship with implicit aftereffects (passive-aftereffects: *p* <.001, *r*^*2*^_*adj*_ =.111, BF_10_ = 34.473; prediction-aftereffects: *p* =.004, *r*^*2*^_*adj*_ =.079, BF_10_ = 7.309). Moreover, a multiple regression with both variables as predictors and angular reach deviations in without-strategy no-cursor trials as the dependent variable, shows that both passive localization shifts (*β* = −0.430, *p* <.001, *sr*^*2*^ =.204) and updates in predicted sensory consequences (*β* = −0.694, *p* <.001, *sr2* =.171) are significantly associated with reach aftereffects (*r*^*2*^_*adj*_=.276, BF_10_ > 5 · 10^4^). Importantly, both hand localization components are still related to implicit reach aftereffects after accounting for a group effect, showing that these relationships are not spurious (data and analysis available on OSF ^[48]^). Furthermore, given that we calculate afferent and efferent contributions to hand localization as additive (see Methods), the two hand localization components are independent from each other (confirmed by a low collinearity: *vif* = 1.087). Finally, we validate our regression model by comparing predicted and observed values of reach aftereffects (Fig. 7c). We find that model predictions are not perfect, but relatively close to observed values (*r*^*2*^_*adj*_=.285, BF_10_ > 3 · 10^5^). The model is likely incomplete, which would explain this disparity, but we don’t investigate this further. In contrast, explicit learning (i.e., with-strategy minus without-strategy reach deviations) has a weak anti-correlation with efferent components of hand localization, and no relation with afferent components of hand localization shifts (Fig. 7d-7f; predicted and observed explicit learning: *p* =.038, *r*^*2*^_*adj*_ =.037, BF_10_ = 0.911). Thus, afferent and efferent-based components of hand localization shifts are weakly, but independently related with implicit reach aftereffects, hinting that at least two separate processes underlie implicit visuomotor adaptation.

**Fig. 7:**
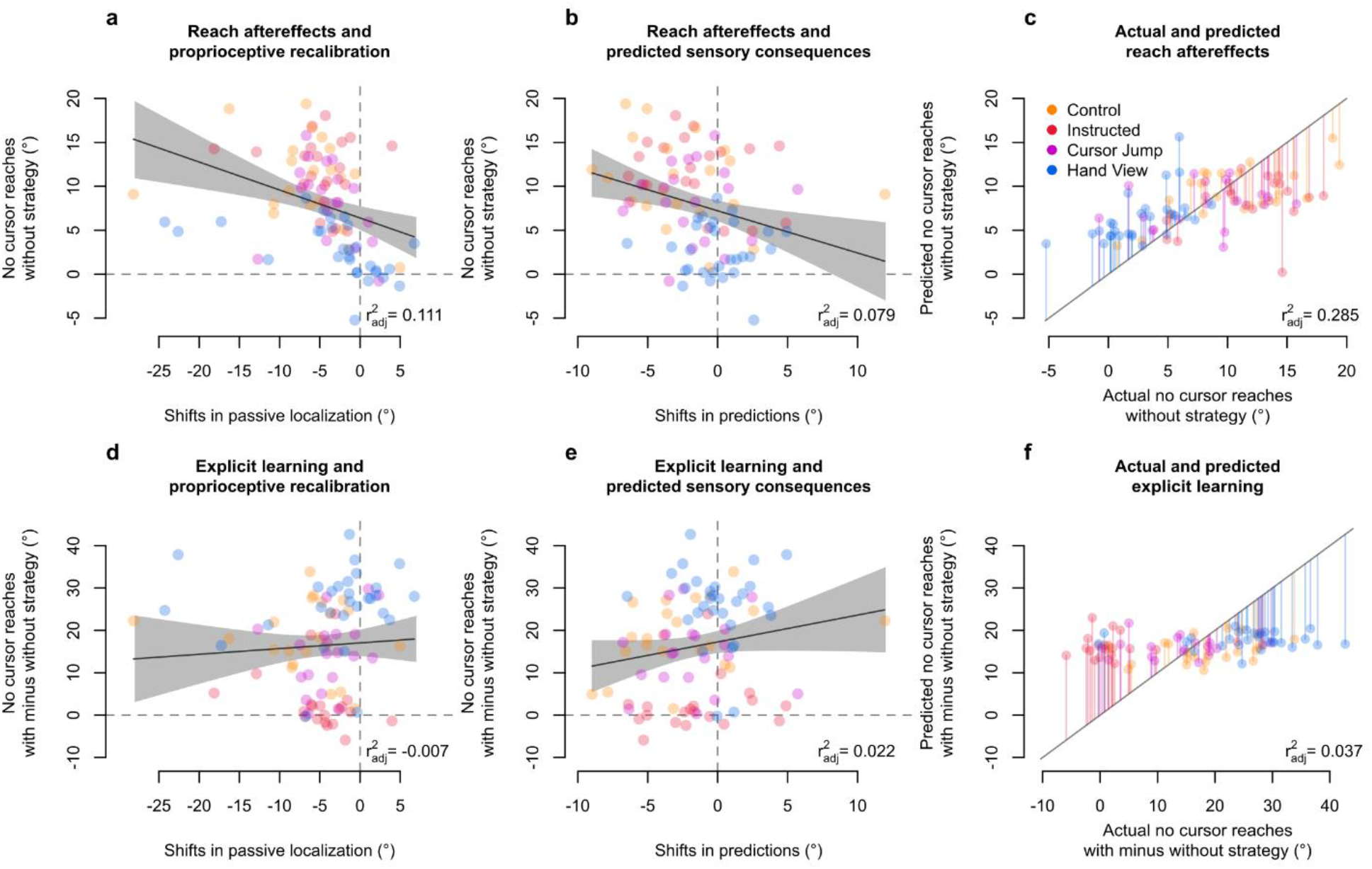
Contributions of afferent and efferent-based hand localization changes to implicit aftereffects and explicit learning. Relationships of afferent and efferent-based changes in hand location estimates with reach deviations when no visual feedback of the cursor is presented, while either excluding any strategies used during adaptation training (implicit aftereffects; **a-b**), or taking the difference of including and excluding such strategies (explicit learning, **d-e**). Individual data points from all participants are colour-coded according to their respective groups. Solid line corresponds to a regression line, while the grey shaded region corresponds to 95% confidence intervals. We then validate the multiple regression model using both shifts in afferent and efferent-based hand localization as predictors, and show the predicted values for reach aftereffects plotted over observed values for reach aftereffects (**c**), as well as the predicted values for explicit learning plotted over observed values for explicit learning (**f**). The diagonal represents perfect prediction. Individual data points are colour-coded according to group, and lines represent residual errors.

## Discussion

We test if manipulating the extent of external error attribution affects both changes in motor behaviour and hand location estimates after visuomotor adaptation training. Particularly, the visual feedback of the hand-cursor either jumps to the imposed rotation mid-reach on every training trial, or is present along with a view of the actual hand of the participant. Given the mismatch between cursor and hand positions, errors should be attributed externally and not lead to changes in hand location estimates. In the Hand View group, despite the error source being clearly external, afferent-based (proprioceptive) hand location estimates still shift to the same extent as in other groups where external error attribution should be minimal. With both afferent and efferent-based estimates (active localization), shifts are not much different in the Hand View group compared to passive localization shifts. Furthermore, we find evidence that the Instructed, Cursor Jump, and Hand View groups developed an explicit strategy. However, the persistent but reduced implicit reach aftereffects in the Cursor Jump and Hand View groups, suggest that the feedback in these groups leads to less implicit learning. The reduction of aftereffects is more profound in the Hand View group, as could be expected with more external error attribution. Finally, we find that both afferent and efferent-based changes in hand localization share a relationship with implicit aftereffects. The persistent implicit motor changes and afferent-based changes in hand position estimates suggest that these are robust against external error attribution, while updating of efferent-based predicted sensory consequences is not.

In visuomotor adaptation, visual feedback of the hand is consistently shifted, which eventually updates estimates of the unseen hand after a movement^[23-24,28,30-32,44-45]^. These updates rely on at least two components: an efferent-based component, where the expected outcome of a motor command is updated to reflect the experienced, altered visual outcome of the movement, and an afferent component, where a proprioceptive signal is recalibrated to the experienced visual outcome ^[29,35,41]^. People should not update either hand location estimate when the rotated cursor does not represent their true hand location. Yet, our previous results suggest that when explicit adaptation increases, due to instructions or increased rotation size, there is no concomitant decrease in updates of afferent and efferent-based estimates ^[18,49]^. In the current study, despite the error source being clearly external in the Hand View group, we surprisingly find shifts in afferent-based hand location estimates across all groups. We also find evidence of efferent-based contributions to hand localization in the other three groups, while this is not so clearly present in the Hand View group. This could mean that heightened external error attribution in the Hand View group decreases efferent contributions to hand location estimates. Nevertheless, proprioceptive recalibration seems to be robust against varying degrees of explicit adaptation and external error attribution.

Changes in afferent-based hand location estimates seem to be a robust form of sensory plasticity, given its relatively quick emergence ^[28-29]^, persistence despite explicit adaptation ^[18]^, and its preservation despite aging ^[30,49]^ and within other forms of perturbations ^[23,31-34]^. Furthermore, recalibrated proprioception is intact in people with mild cerebellar ataxia ^[50]^, despite the cerebellum’s crucial role in adaptation ^[1,3,14,44-46,51-52]^. This implies that proprioceptive recalibration relies on a signal different from efferent-based contributions to hand localization, such as a visuo-proprioceptive discrepancy ^[29,35,41]^. Although there should be no visuo-proprioceptive discrepancy in the Hand View group, as they see and feel their hand, our findings suggest otherwise. Since the task is completed by bringing the cursor to the target, the cursor could be acting as a visual placeholder for the actual hand, while proprioceptive feedback is still veridical. This could create a sensory discrepancy, between seen cursor and felt hand, leading to sensory recalibration. Thus, the Hand View group does not show decreased shifts in afferent-based hand localization, despite external error attribution. It also seems that in only the Hand View group, there might not be an efferent-based contribution to hand localization, or one that is hard to detect. While this will have to be replicated, it is in line with previous findings ^[35,41]^ that also indicate that efferent and afferent contributions to hand localization rely on different error signals.

Aside from sensory recalibration, visuomotor adaptation also leads to implicit motor behaviour changes. Implicit learning is rather stable, but awareness of the perturbation’s nature increases explicit contributions during adaptation^[15-16,18-22,53-54]^. Here, participants make open-loop reaches with (implicit and explicit) or without (implicit) the strategy they learned. This process dissociation procedure (PDP, ^[16]^) is consistent with similar tasks ^[53,55]^, has been used in previous studies ^[17-18,43,56]^, and doesn’t seem to evoke additional explicit learning unlike other methods ^[56-58]^. While explicit learning does not necessarily correspond to external error attribution, it is likely that external error attribution is accompanied by more explicit adaptation. Despite no elaborate instructions, the Cursor Jump and Hand View groups exhibit explicit learning like the Instructed group. Furthermore, it seems advantageous to suppress implicit learning with external and likely transient perturbations ^[6-10]^, making adaptation largely explicit or strategy-based ^[59-60]^. Here, although implicit learning persists, we observe a small decrease in implicit adaptation in the Cursor Jump group, which is much more pronounced in the Hand View group. Although we expect increased external error attribution in the Cursor Jump and Hand View groups, this effect seems to be less clear for the Cursor Jump group. Nonetheless, we are certain that the Hand View group attributes the source of the error more externally than other groups.

A reduction of sensory prediction error-based learning may explain the reduced reach aftereffects and efferent-based hand localization shifts in the Hand View group. Implicit adaptation is based on sensory prediction errors ^[3-4,13,44-46]^, that both healthy individuals and people with cerebellar damage involuntarily engage in^[1,3,14,46,52]^. In the Hand View group, the balance between sensory prediction error-based learning and explicit strategy contributions to behaviour is changed. Consistent with previous studies using a similar condition as the Hand View group ^[46,59-60]^, our data suggest that increased external error attribution leads to reduced sensory prediction error-based visuomotor adaptation. Furthermore, efferent-based updates in predicted sensory consequences contribute to hand location estimates. The decreased sensory prediction error-based learning should result in little to no shift in efferent-based hand position estimates. Thus, while afferent-based contributions to hand localization rely on visuo-proprioceptive discrepancy signals, changes in efferent-based contributions depend on sensory prediction error-based learning. Consequently, it seems that external error attribution only reduces sensory prediction error-based learning.

Reach aftereffects are evidence that people have updated their internal model, and hence efferent-based predictions, to adapt movements ^[5,11-12]^. Recalibrated proprioception also informs movements ^[29,35-37,47,50,61]^. First, preventing updates of internal models while allowing for proprioceptive recalibration, leads to reach aftereffects that follow the proprioceptive shift ^[29,32,35-37,47,50]^. Second, recalibrated proprioception is at maximum within six trials or faster ^[28-29]^. Both these findings make it unlikely that proprioceptive recalibration arises due to repeated hand movements performed during adaptation. One likely interpretation is that both changes in efferent-based predictions and recalibrated proprioception separately contribute to changes in motor behaviour (reach aftereffects). Here, we show with a multiple regression that both afferent and efferent changes are independently related to reach aftereffects in without-strategy no-cursor reaches. Given that, for now, we consider afferent and efferent contributions as additive in hand localization (see Methods), these contributions are necessarily statistically independent from each other. Moreover, our behavioural evidence shows that suppressed efferent-based changes in the Hand View group are tied to reduced implicit reach aftereffects. Based on these results, we speculate that the remaining reach aftereffects for the Hand View group are solely based on afferent changes. Regardless, our data show that changes in motor behaviour after learning take into account updates to our multi-modal internal estimates of hand location.

The changes in both afferent and efferent-based hand location estimates that rely on different error signals, and are independently related with changes in motor behaviour, are likely processed in different regions of the brain. While the relationship between implicit adaptation and sensory prediction error-based learning has been linked to the cerebellum ^[3-4,13,44-46]^, the visuo-proprioceptive discrepancy leading to recalibrated proprioception has been linked to parietal areas ^[25,31,62-63]^. Particularly, parietal lesions that disrupt the angular gyrus in the posterior parietal cortex (PPC) affect the relationship between the weighting of visuo-proprioceptive information and corresponding realignment ^[62]^, which in turn affects corresponding activity in somatosensory and motor areas ^[25,63]^. In the current study, the greatly reduced efferent-based changes and persistent afferent-based changes in hand location estimates, due to external error attribution in the Hand View group, show that processing for these two signals is dissociated to some degree in the brain. However, although afferent and efferent-based signals seem to be independently processed in brain, both the PPC and cerebellum have connections with premotor and motor cortical areas ^[25,63]^. Here, we do find that afferent and efferent-based hand location estimates share small but significant relationships with implicit reach aftereffects. Thus, our data are consistent with the interpretation that the independent signals used in updating our hand location estimates are likely integrated within premotor and motor areas, and consequently affect our motor behaviours.

In summary, external error attribution affects changes in our internal estimates of hand location and motor behaviour. Particularly, changes in afferent-based (proprioceptive) estimates of hand location are so robust, that the resulting recalibration is unaffected by external error attribution. However, external error attribution can be manipulated to change efferent-based, sensory prediction error-based learning. As adaptation becomes less reliant on sensory prediction error-based learning, implicit motor behaviour changes (reach aftereffects) are consequently reduced. We also find behavioural evidence that these afferent and efferent-based estimates contribute independently to motor behaviour changes. Taken together, it seems that proprioceptive plasticity plays an important role when updating our hand location estimates after experiencing movement errors, as sensory prediction error-based processes are reduced with increased external error attribution, but visuo-proprioceptive recalibration is impervious to this.

## Methods

### Participants

Ninety right-handed university students (64 female, *M*Age = 20.8, *SD*_Age_ = 3.88) were assigned to one of four groups: Control (*n* = 20, 14 females), Instructed (*n* = 21, 13 females), Cursor Jump (*n* = 20, 14 females), and Hand View (*n* = 29, 23 females). Data for the Instructed and Control groups have been used in our earlier work and are publicly available on OSF ^[18]^. In those two data sets, the samples (∼20 participants per group) were large enough to detect differences between active and passive localization shifts (see also ^[41]^, *n* = 19). For the Cursor Jump group, the sample size matched these reference groups. Since, to our knowledge, no previous study has compared active and passive hand localization shifts after training with a full view of the hand, we ensured sufficient power to detect subtler effects by adding more participants to the Hand View group. All participants gave written informed consent prior to participating. All procedures were in accordance with institutional and international guidelines. All procedures were approved by York University’s Human Participants Review Committee.

### Experimental Set-up

#### Apparatus

Participants held the handle of a 2-joint robot manipulandum (Interactive Motion Technologies Inc., Cambridge, MA, USA) with their right hand, while placing their thumb on top of the handle. A downward facing monitor (Samsung 510 N, 60 Hz) 28 cm above the manipulandum projected visual stimuli on a reflective tint (14 cm above the manipulandum), making the stimuli appear on the same horizontal plane as the participant’s hand (Fig. 1a-1c). The reflective tint is applied to plexiglass and achieves the same result as a half-silvered mirror. Participants responded using their visible left hand in some tasks on a touchscreen 2 cm above the manipulandum (Fig. 1d). The right hand was occluded from the participant’s view and a black cloth was draped over their right arm and shoulder. For the Hand View group, the right hand was illuminated in some tasks, making it visible to the participant.

#### Stimuli

Participants made smooth and straight 12 cm out-and-back reaching movements from the “home position” to one of three targets (or arcs). Targets and arcs were presented once in a shuffled order before being presented again, such that reach directions were evenly distributed across trial types (Fig. 2).

#### Cursor Training Trials

Participants kept a green cursor (circle, 1 cm diameter), representing their right thumb, at the home position for 300 ms. A yellow target (circle, 1 cm diameter) then appeared at one of three possible locations: 45°, 90°, 135° in polar coordinates. Once the target was acquired, they held the cursor for 300 ms within 0.5 cm of the target’s centre. Afterwards, both stimuli disappeared, and participants returned their hand to the home position via a robot-constrained path (perpendicular resistance force: 2 N/(mm/s); viscous damping: 5 N/(mm/s)). Participants in the Hand View group saw their right hand along with the cursor throughout these trials. For these trials, we calculated the angular difference between the hand position at the peak of movement velocity and the target, relative to the home position. Thus, once the rotation is introduced, full adaptation should then result in angular reach deviations of 30°.

#### No-Cursor Trials

These proceeded similarly to cursor training trials, but without visual feedback from the cursor or hand (Fig. 1f). Participants kept stationary for 500 ms once they believe they had acquired the target with their unseen right hand, making the target disappear. They returned to the home position via the constrained path.

During the rotated session, participants completed two variations of no-cursor trials in succession (with- and without-strategy; Fig. 2). Using the process dissociation procedure from Werner et al. (PDP; ^[16]^), we instructed participants to either include or exclude any consciously accessible strategy they developed to counter for the visuomotor rotation, to measure implicit and explicit adaptation. The order of these blocks was counter-balanced within one participant and between participants (Fig. 2). For all no-cursor trials, we calculated the angular difference between the endpoint of the participant’s hand movement and the target, relative to the home position. Considering reach endpoints makes this data set comparable to those from localization trials.

#### Localization Trials

Participants saw a white arc (0.5 cm thick) 12 cm away from the home position (Fig. 1e), which spanned 60°, and was centred on either the 50°, 90°, or 130° mark in polar coordinates. In self-generated active localization trials, participants moved their unseen right hand from the home position to any point on the arc, and were instructed to vary their chosen crossing points across trials. In passive localization trials, the robot guided the participant’s right hand towards the same points on the arc that they intersected during active localization trials in the preceding task. Regardless of localization type, a cushion force prevented hand movements from moving beyond the arc position. Participants then voluntarily returned their right hand to the home position via the constrained path, and used their visible left hand to indicate on the touchscreen the point at which they believed their unseen right hand intersected the arc.

#### Procedure

The aligned session served as baseline data, and started with aligned cursor training trials, followed by blocks of active localization, passive localization, and no-cursor trials respectively (Fig. 2). Localization and no-cursor blocks were repeated in the same order for three more times during this session. To prevent decay in learning, we interleaved shorter blocks of “top-up” cursor training trials between every localization and no-cursor block. The aligned session ended upon completion of the fourth no-cursor block.

Participants were given a mandatory five-minute break. During this break, the Instructed group was informed about the nature of the perturbation and was given a strategy to counter it (see ^[15,18]^ for details). The other groups were simply advised to compensate since the cursor would be “moving differently”, and to remember any strategy they develop as they would be asked to either use or not use it.

In the following session, the cursor was rotated 30° clockwise (CW) relative to the hand position for all cursor training trials. Hence, correcting for this perturbation requires straight reaches in the 30° counterclockwise (CCW) direction. Regardless of instructions received during the break, both Instructed and Control groups simply experienced this perturbation. For the Cursor Jump group, the cursor shifted to this rotated trajectory after participants moved for one-third (4 cm) of the home-target distance (Fig. 1b). For the Hand View group, illuminating the right hand allowed participants to see the misalignment between cursor and hand, making this the clearest demonstration that the error was caused externally (Fig. 1c). The rotated session proceeded similarly to the aligned session. However, to saturate learning of the visuomotor rotation, we increased the number of cursor training trials in each block (Fig. 2). Moreover, each block of no-cursor trials was done twice, each in one variation (with-strategy or without-strategy).

## Data Analysis

We compared all four groups within the different trial types. Results from frequentist tests are reported with an alpha level of 0.05. Greenhouse-Geisser corrections were applied when necessary. Planned follow-up tests used the Sidak method when it was necessary to correct for multiplicity. Degrees of freedom for follow-up tests are larger than expected in some cases, as it uses a model fit on all the data (R emmeans package, ^[64]^). For the figures, estimates of confidence intervals were bootstrapped to represent the individual data better, but confidence intervals for grouped data and the corresponding statistical tests were based on sample t-distributions. All data preprocessing and analyses were conducted in R version 3.6.0^[65]^. Bayesian statistics are reported for each corresponding frequentist test and were conducted in JASP version 0.11.1 ^[66]^. Follow-up tests for Bayesian ANOVAs were only conducted on main effects (*odds* values in Results). We conducted Bayesian t-tests to follow-up on interaction effects, without correcting for multiplicity.

### Rate of Learning During Adaptation Training

We analyzed cursor training trials from both the aligned and rotated sessions. Trials were manually inspected for outlier reaches (0.94% of trials removed). We corrected for individual baseline biases by calculating the average reach deviation for each target separately within each participant, during the last 30 out of the first 45 aligned cursor training trials, and subtracting this from rotated cursor training trials. We compared angular reach deviation measures across all groups, within each one of three trial sets (rotated cursor training trials 1-3, 4-6, 76-90), to confirm learning and investigate any differences.

### Reach Aftereffects and Strategy Use

We tested for group differences in reaches without cursor-feedback. Upon manual inspection, outlier reaches were removed (1.46% of trials). We confirmed the presence of reach aftereffects by comparing angular reach deviations from aligned no-cursor trials to without-strategy no-cursor trials. For the PDP ^[16,18]^, we implemented baseline-correction (aligned session no-cursor reaches subtracted from no-cursor with- and without-strategy trials, respectively), before comparing angular reach deviations in with- and without-strategy trials.

### Proprioceptive Recalibration and Updating Predicted Sensory Consequences

We investigated active and passive localization trials, before and after adaptation training. We calculated the angular difference between the endpoint of the participant’s right hand movement and their left hand responses on the touchscreen, relative to the home position. Localization response biases were accounted for using a circle fitting procedure (see ^[35]^ for details). Trials with hand movement endpoints beyond ±20° from the arc centre across all groups, and angular errors beyond ±3 standard deviations from the mean angular error per participant were removed (1.06% of angular errors). We used a kernel smoothing method (gaussian kernel with bandwidth = 15°) to interpolate changes in hand localization at specific points (50°, 90°, 130°) for every participant. Mean values at each of these points estimate active and passive hand localization errors for both the aligned and rotated sessions.

We compared hand localization errors in the rotated session to those in the aligned session. The difference of localization errors between the two sessions represents shifts in hand localization, and were compared across groups and movement type (active and passive). The difference between active and passive localization shifts were used as a measure of efferent-based updates in predicted sensory consequences, while passive localization shifts measured the afferent-based recalibration of proprioception. If afferent and efferent contributions to hand localization are optimally integrated (e.g. Bayesian integration), then variance in active localization should be lower than passive localization ^[41]^. However, we have failed to find this in two earlier studies^[35,41]^ as well as more recently, when we combined data from several studies, for a total of over 200 participants ^[67]^. Thus, we take a parsimonious approach, and treat afferent and efferent contributions as additive in hand localization. We compared these measures for each group against zero, and investigated how both hand location estimates may contribute to implicit motor changes.

## Data Availability

Data, analyses scripts, and preprint are available on Open Science Framework (https://doi.org/10.17605/osf.io/xdgh6 ^[48]^).

## Acknowledgements

This work was supported by NSERC for DYPH; SSHRC, OGS, and VISTA for RQG; OGS, and VISTA for SM. The funders had no role in study design, data collection and analysis, decision to publish, or preparation of the manuscript.

## Author Contributions

BMtH and DYPH designed the research. RQG and SM collected the data. BMtH contributed experimental and analytic code. RQG, SM, and BMtH analyzed the data. RQG wrote the manuscript, which was carefully edited by all authors. The final version of the manuscript has been approved by all authors who agree to be accountable for all aspects of the work in ensuring that questions related to the accuracy or integrity of any part of the work are appropriately investigated and resolved.

## Conflict of Interest

The authors declare no competing interests.

## Notes

https://doi.org/10.17605/osf.io/xdgh6

### Competing Interest Statement

The authors have declared no competing interest.

### Summary of Updates

Bayesian statistics added; clarifications to definition of concepts and methods used; Figure 7 updated

